# The effects of conditional loss of myosin binding protein H-like on cardiac function

**DOI:** 10.1101/2024.10.07.617136

**Authors:** Hope V. Burnham, Alex Peña, Shreeya Gandhi, Geena Fritzmann, David Y Barefield

**Author notes:** To whom correspondence should be addressed, David Y. Barefield, PhD, Department of Cell and Molecular Physiology, Loyola University Chicago, 2160 S. 1st Ave, Maywood, IL 60153.

## Abstract

Mutations in the myosin-binding protein H-like (MyBP-HL) gene, *MYBPHL*, are linked to hereditary dilated cardiomyopathy (DCM), atrial fibrillation, and atrioventricular arrhythmias. MyBP-HL is a sarcomeric protein that is highly expressed in the atria with only scarce, distinct clusters of MyBP-HL positive cells within and surrounding the ventricular conduction system. Constitutive knock-out of MyBP-HL in mice causes atrial dilation, arrhythmia, and DCM. Whether MyBP-HL plays a developmental role, or if knock-down in adulthood will recapitulate a similar phenotype has yet to be examined. Moreover, the significance of the MyBP-HL expressing ventricular cells, or the functional need for differential thick filament regulation is currently unknown.

We used a conditional floxed *Mybphl* mouse to further elucidate the role of MyBP-HL. We crossed this mouse with a ROSA26-Cre(ERT2) LoxP mouse to conditionally knock-down *Mybphl* after tamoxifen treatment. We also crossed the *Mybphl* flox mouse with a Contactin-2-Cre mouse that deletes *Mybphl* solely in the cardiac conduction system from birth. Echocardiography was used to measure contractile function, and conscious telemetry allowed for monitoring of heart rhythm and electrical signal conduction changes.

We demonstrate that mice with conditional decrease of MyBP-HL in adulthood develop a hypertrophic phenotype with atrial contractile changes, increased total heart weight to body weight, and increased heart rate variability. Deletion of *Mybphl* solely within the cardiac conduction system trends toward mild hypercontractility, lower heart rates, and interventricular septal thickening. These data show that MyBP-HL is essential for proper cardiac function, and even minor alteration in protein levels cause a diseased cardiac phenotype.

## INTRODUCTION

The discovery of the sarcomere protein myosin binding protein H-like (MyBP-HL) has many ramifications for our scientific understanding of thick filament regulation. Whereas previously, it was thought that the heart had only one main myosin binding protein, MyBP-HL is highly expressed in the atria and is found in a subset of cardiomyocytes around the ventricular conduction system (VCS) [1, 2]. Characterization of this protein in recent years demonstrates a role for the dysregulation of MyBP-HL in the onset of human cardiac disease, linking it to both dilated cardiomyopathy (DCM), atrial fibrillation (AF), and atrioventricular arrhythmias [1, 2]. Constitutive null (*Mybphl^-/-^*) and heterozygous (*Mybphl^WT/-^*) mouse models recapitulate the DCM phenotype, with increased left ventricular internal diameter, atrial dilation, and an increased rate of arrhythmias [1]. Single atrial myofibrils lacking

MyBP-HL demonstrate decreased linear relaxation time and an increased rate of relaxation compared to wild type atrial myofibrils, implicating MyBP-HL for a potential role in slowing cross-bridge cycling [3]. MyBP-HL expression is seen in all atrial cardiomyocytes, and it is known to be expressed very early in development, at least at embryonic day 9.5 [4, 5]. In fact, the levels of the related protein, cardiac myosin binding protein-C (cMyBP-C) in the atria are half that measured in the ventricles [3]. Moreover, protein analysis via immunoblotting and mass spectroscopy demonstrate competition between MyBP-HL and cMyBP-C for thick-filament binding sites. It is known that per half-sarcomere, there are 27 sites on the thick filament for myosin binding proteins to associate. An inverse relationship between MyBP-HL and cMyBP-C exists, wherein healthy sarcomeres have a 50/50 split of both myosin binding proteins bound to myosin, yet decreased expression of one protein causes an increased amount of the other protein to be found in sarcomeres and *vice versa* [3]. The effect of MyBP-HL has almost exclusively been studied in the context of constitutive loss of the protein. Whether conditional loss of MyBP-HL in a fully formed, adult, healthy heart would cause disease, or if there is a developmental window in which MyBP-HL is required for normal development, is unclear.

Beyond its high expression in atrial cardiomyocytes, MyBP-HL is also found in a select number of ventricular cardiomyocytes that appear to preferentially express in or around the ventricular conduction system (VCS) [2]. With the observation of arrhythmias and atrioventricular conduction abnormalities in human patients with MyBP-HL mutations, including multiple cases of complete heart block, and murine model arrhythmias, a potential role for MyBP-HL in cardiac conduction is proposed [1, 2]. The unique expression of distinct MyBP-HL foci associated with the VCS further supports a need to investigate whether loss of these localized regions of MyBP-HL expression specifically in the VCS would be consequential.

To answer these questions, we used two Cre mouse models crossed with a *Mybphl*^(flox/flox)^ mouse. The first is a Rosa26-Cre(ERT2) model that recombines and effectively suppresses the *Mybphl*^(flox/flox)^ gene after administration of tamoxifen throughout the body. The model allows temporal control of gene knock-down and is used to knock-down MyBP-HL in adulthood upon tamoxifen administration around 11-12 weeks of age. Functional analysis before and after tamoxifen administration demonstrates whether loss of MyBP-HL in adulthood still causes disease, identifying if MyBP-HL is essential for healthy functioning in the human heart. The other mouse model used in this project is a contactin-2-Cre mouse, wherein the *Mybphl*^(flox/flox)^ gene is recombined solely in cells expressing the contactin-2 gene, which in the heart is a marker of the conduction system. This model provides a way to determine possible roles for the specific subset of MyBP-HL positive cells within the conduction system of the heart.

## MATERIALS AND METHODS

### Animal Models

A mouse line (*Mybphl^tm1c(KOMP)Wtsi^*) with LoxP sites flanking exons 2 - 6 of the *Mybphl* gene was obtained through the Knock Out Mouse Project [6]. These flanking LoxP site (“floxed”) *Mybphl*^(flox/flox)^ mice were bred with either Rosa26-Cre(ERT2) mice (Jackson *B6.129-Gt(ROSA)26Sor^tm1(cre/ERT2)Tyj^/J* stock #008463) [7] or Contactin-2-Cre *(Cntn2^3’UTR-IRES-Cre-EGFP/+^)* mice [8]. Both lines were maintained on a C57Bl6/J background and housed in a specific pathogen-free, environmentally controlled facility. The Cre alleles for both mouse lines were maintained in a hemizygous state, providing Cre-negative and Cre-positive littermates. All animal studies and procedures were approved by the Loyola University Chicago Institutional Animal Care and Use Committee, and animals were housed and treated with the standards set by the PHS Policy on Humane Care and Use of Laboratory Animals. Male and female subgroups were used for all experiments.

### Tamoxifen Treatment

Rosa26-Cre(ERT2) mice were used starting at 11- to 15-weeks of age, as specified. Mice were treated with 100 μL of 25 mg/mL tamoxifen (Sigma Aldrich T5648-1G) that was first diluted in corn oil (Sigma C8267) and passed through a sterile 0.2-μm filter. Tamoxifen was administered via intraperitoneal injection for 4 consecutive days. Mice were analyzed 6 weeks post-injection.

### Echocardiography

Mice were anesthetized using 5% vaporized isoflurane in 100% O_2_ during induction and then 1% isoflurane at 1 L/min for the plane of anesthesia. Mice were placed in the supine position onto a heated table and surface ECG recordings were taken for monitoring. Fur was removed from their chests using topical hair removal cream. Echocardiography was performed using an MS550D 22- to 55-MHz solid-state transducer on a Visual Sonics Vevo 3100 imaging system (FujiFilm, Toronto, Canada). The following views were recorded: Parasternal long axis, parasternal short axis (at mid-papillary level), apical four-chamber using color doppler, pulse wave doppler, tissue doppler, and a left lateral long-axis for 4D EKV recordings of the left atrium. Images were analyzed using the Vevo Lab software. Both acquisition and analysis were conducted blinded to experimental conditions.

### Cardiac Telemetry

Wireless implantable cardiac telemetry transmitters (TSE Systems, E-430001-IMP-62) were subcutaneously implanted surgically in mice under isoflurane anesthesia. The mice were allowed to recover for one week prior to data collection. Overnight electrocardiography recordings were taken to establish baseline characteristics. On subsequent days, mice were injected with isoproterenol (4 mg/kg), propranolol (1 mg/kg), and flecainide (20 mg/kg) [1, 2]. Telemetry data was recorded for 45 min following drug administration. ECG recordings were analyzed using the Notocord-hem analysis software with the QRS10a and ECG51a modules. For each mouse and condition, a 10 min segment of telemetry data was chosen for in-depth analysis. Analysis was performed blinded to experimental conditions.

### Tissue Collection

Mice were humanely euthanized in accordance with the IACUC protocol then dissected to remove the hearts. The hearts were rinsed with cold 1x PBS and the atria were isolated. Body weight, total heart weight, and atrial weight were measured at the time of collection. Tissue samples were flash-frozen using liquid nitrogen and stored at −80° C until use. For the Rosa-26-Cre(ERT2) mice, liver samples were also collected and flash frozen and tail clippings were collected and stored at 4° C for DNA digestion. For the Contactin-2-Cre mice, in addition to the atria being isolated, hearts were cut open coronally and samples were collected corresponding to the atrioventricular (AV) node, left ventricular endocardial wall, right ventricle, and non-endocardial left ventricle. These were also flash frozen and stored at −80° C.

### DNA Isolation

Approximately 5-10 mg per tissue had 300 μL of cold cell lysis solution (Quiagen #158908) with 1.5 μL of 20 mg/mL proteinase K solution (Quiagen #19131) added. Samples were incubated overnight at 55° C then cooled to room temperature. Samples were vortexed thoroughly after addition of 100 μL protein precipitation solution (Quiagen #158912), and centrifugation was performed at 16,000 x g for 3 min to pellet the protein. The DNA was precipitated by adding 300 μL of isopropanol to the supernatant. The samples were mixed thoroughly by inversion then centrifuged at 16,000 x g for 1 min. The supernatant was discarded, and the DNA pellet was rinsed using 300 μL of 70% ethanol via inversion. Samples were centrifuged at 16,000 x g for 1 min, supernatant discarded, then sample tubes were inverted and allowed to air dry for 15 min. DNA pellets were resuspended in 30 μL sterile DNase/RNase free water, and allowed to rehydrate overnight at RT. Samples were stored at −20° C prior to use.

### Polymerase Chain Reaction and Gel Electrophoresis

Polymerase Chain Reaction (PCR) was performed in 10 μL final volumes with 0.05 Units of DreamTaq and DreamTaq buffer (Thermo Fisher EP0703), 0.2 mM dNTPs (Thermo Fisher FERR0241), 10 μM of each primer, and 1 μL DNA. Tail and cardiac DNA samples were diluted 1:10, and liver samples were diluted 1:100 in sterile water prior to addition to PCR tubes. Primers flanking and within the LoxP sites were designed for the unrecombined and recombined alleles (forward 1: 5’-GGC GCA TAA CGA TAC CAC GA-3’; forward 2: 5’-GTT ACT TAG GCA CTG TCA GGA AGG G-3’; reverse: 5’-GCA GAG CAG TAT GCA TGT CCA C-3’). The samples were amplified in the BioRad C1000 Touch Thermo Cycler using the following cycle conditions: 94° C x 2 min, (94° C x 15 sec, 63° C x 30 sec, 72° C x 30 sec) x35, 72° C x 10 min. Samples were run mixed with Tri-Track Loading Dye (Fisher Scientific FERR1161) on a 2.5% agarose gel with ethidium bromide. Gel was then imaged in an Azure 200 BioSystems Gel Imager.

### Protein Isolation

Tissue samples were homogenized in TriZOL (Fisher Scientific 15596018) and then were cooled and separated into aqueous and organic layers using chloroform (Millipore Sigma C2432). The aqueous layers were carefully pipetted off for RNA isolation. Any remaining aqueous phase in the tubes was pipetted out, then 300 μL of 100% ethanol per 1 mL TriZOL used for initial homogenization was added to the TriZOL-chloroform layer. Samples were mixed by inversion and allowed to incubate for 2-3 min at RT. Samples were centrifuged at 2,000 x g for 5 min at 4° C to pellet the DNA, then supernatant was collected. Between 200 and 400 μL of supernatant was used to isolate protein. Any remaining supernatant was stored at −20° C. Protein isolation was then performed according to Thermo Fisher Protein Isolation from TriZOL protocol (Thermo Fisher AM9738).

The protein pellets were re-solubilized in a urea buffer (8 M Urea (Thermo Fisher U15-500) with 0.3% TritonX-100 (Millipore Sigma X100-100mL) and Protease Inhibitor (Thermo Fisher A32955)) at approximately 100 μL buffer per 5 mg of starting tissue. The pellets were gently dispersed and solubilized in solution for 10-15 min by periodically “flicking” the tube. The samples were heated to 100° C for 3 min in a heat block, centrifuged at 10,000 x g for 5 min to pellet insoluble material, and the supernatant was transferred to a new tube. Protein concentrations were determined using the BioRad Bradford Reagent kit (5000205). Samples were stored at −20° C until use.

### Immunoblotting

Protein sample aliquots were standardized and diluted using the urea buffer solution and 2x Laemmli Buffer (BioRad 1610737) to be used for immunoblotting. Total protein lysate samples were run on 10% acrylamide gels with a Chameleon ladder (LI-COR 928-60000). Prior to loading, samples were heated at 75-80° C for 5-10 min and vortexed thoroughly. Gel electrophoresis was run at 130 V for approximately 1 hour in tris glycine-SDS running buffer (25 mM Trizma base, 190 mM Glycine (Fisher Scientific AAA138160E), and 0.1% SDS). The gel was transferred to a PVDF membrane at 300 mA for 3 hrs in cold Tris-Glycine Methanol (25 mM Trizma base, 190 mM Glycine, 20% (V:V) methanol (Thermo Fisher 041467.K7)) solution.

Prior to blocking the membrane, total protein staining was performed using the LI-COR Revert 700 Total Protein Stain Kit (LI-COR 926-11010), according to kit instructions. The membrane was blocked with 1% casein blocker (BioRad 1610782) for 30-60 min. The membrane was incubated overnight in blocking buffer with primary antibodies: MyBP-HL 1:2500 [2]; cMyBP-C 1:2500 (Santa Cruz 137180), Contactin-2 1:1000 (R&D Systems AF4439), MLC2v 1:2000 (Proteintech 10906-1-AP), and MLC2a 1:2000 (SYSY Antibodies 311 011). Secondary antibodies were IRDye DaM 680rd 1:5000 (LI-COR 926-68072); IRDye GaR 800cw 1:5000 (LI-COR 926-32211) and imaged using Azure Biosystems Sapphire Biomolecular Imager. All blots were analyzed via densitometry in Azure Spot Pro.

### Tissue Clearing

Adult mouse hearts underwent tissue clearing using the iDISCO+ protocol as described previously [2]. Steps were performed at room temperature unless stated otherwise. Briefly, mouse hearts were cannulated and perfused with PBS followed by 4% PFA in PBS. Samples were fixed in 4% PFA in PBS overnight at 4° C with shaking then for 1 hr at room temperature the following morning. The samples were rinsed in PBS with shaking for 30 min x3. Samples underwent a dehydration series, hydrogen peroxide bleaching, then rehydration. To prepare samples for immunostaining they were permeabilized and then blocked. Primary then secondary antibody solution steps were performed at 1:500 concentrations. The primary antibodies were MyBP-HL [2] and Contactin-2 (R&D Systems AF4439). Secondaries were Alexa Fluor Donkey anti-Rabbit 568 (Thermo Fisher A-10042) and Alexa Fluor Donkey anti-Goat 647 (Thermo Fisher A-21447). To clear the tissue, samples were again dehydrated in a methanol/H_2_O series then incubated for 3 hrs in 66% dichloromethane and 33% methanol on a rocker. Two 15-minute rinses were performed with 100% dichloromethane to remove residual methanol. Samples were stored in dibenzyl ether and stored at 4° C until imaging.

### Immunofluorescence Microscopy

Samples that underwent iDISCO+ tissue-clearing protocol were imaged using a Zeiss Light Sheet Microscope 880 with Airyscan microscope. The tissue samples were immersed in dibenzyl ether in a 35 mm MatTek dish. Tiled Z-stacks were acquired and stitched together to create a 3D rendering of the tissue based on fluorescent signal. Files were analyzed using Imaris Imaging Software (Oxford Instruments) to render the MyBP-HL signal as spots and the Contactin-2 signal as a surface. The distance between spots and surface was measured. A tissue sample that was solely incubated with secondaries was used to set parameters for quantification to minimize off-target secondary signal.

### Statistics

Statistical analyses were performed using Prism Version 9.2.0 (GraphPad Software, La Jolla, California). Data were tested for normality using the Shapiro-Wilk test. If data fit a normal distribution, significance was determined using Student’s t-test, one-way analysis of variance (ANOVA), or two-way repeated measures ANOVA where stated. If data did not fit normal distribution, analysis for outliers was performed and/or data was analyzed using non-parametric tests such as Mann-Whitney and Kruskal-Wallis, where applicable. Significance was determined as p ≤ 0.05. Data are reported as individual values overlying mean ± SEM.

## RESULTS

### Tamoxifen Treatment Recombines the Mybphl Allele and Reduces Protein Level in Rosa-26-Cre(ERT2) mice

To confirm recombination of the *Mybphl*^(flox/flox)^ allele in multiple tissue types, DNA was isolated from tail clippings taken before tamoxifen injections began and then approximately eight weeks following injections at time of tissue collection **(Figure 1A)**. DNA was additionally isolated from ventricular and liver tissue. A three-primer assay was designed using two different forward primers to amplify two products, one being the unrecombined *Mybphl*^(flox/flox)^ allele and the other specific to the recombined allele, as shown in **Figure 1B**.

**Figure 1.**
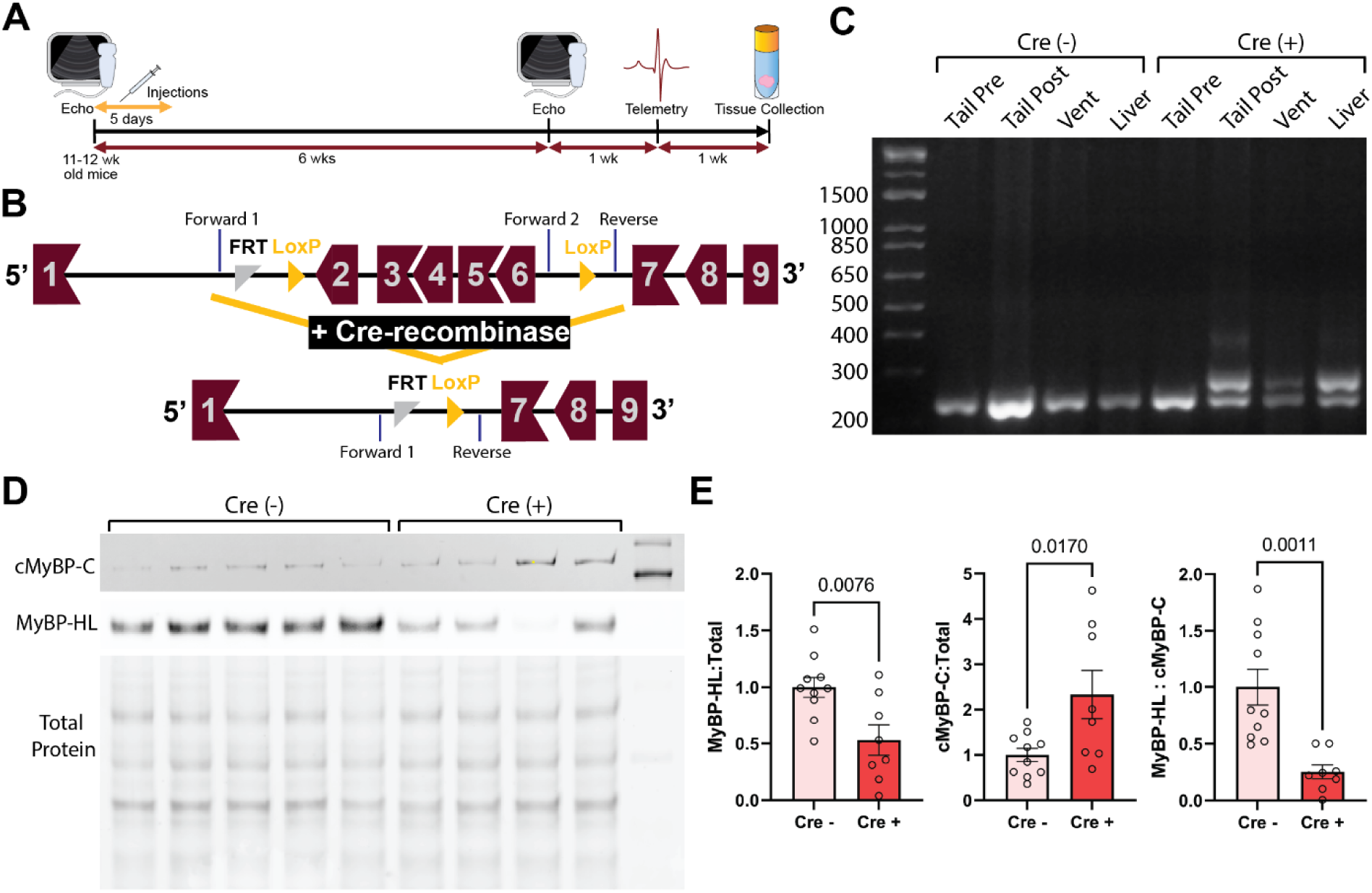
Knock-Down of *Mybphl* in Adult Mice. **A)** Timeline for tamoxifen injections, knockdown period, and physiological measurements. **B)** LoxP sites flank *Mybphl* exons 2 through 6, removing the central exons and altering the reading frame, resulting in a null allele. PCR primer locations before and after recombination are shown. **C)** Agarose gel showing genomic PCR products in the following mouse tissue samples: Pre-tamoxifen tail clipping (Tail Pre), and post-tamoxifen tail clipping (Tail Post), ventricle (Vent), and liver. Samples are grouped together by mouse. **D)** A representative immunoblot of protein lysates from Cre-negative (n = 5) and Cre-positive (n = 4) atria showing levels of MyBP-HL and cMyBP-C. **E)** Ratio of MyBP-HL to total protein, cMyBP-C to total protein, and MyBP-HL to cMyBP-C in atrial tissue lysates. Cre-negative: n = 10, Cre-positive: n = 8. Analysis performed by Student’s T-test.

The unrecombined gene amplicon is 233 bp in length, whereas the higher band, 270 bp, is the recombined gene product. The Cre-negative mice only show the single band at 233 bp across all tissue types, whereas the Cre-positive mice have recombination bands after tamoxifen in the tail, ventricle, and liver samples, demonstrating effective recombination **(Figure 1C)**. The presence of the unrecombined band that persists after recombination indicates that tamoxifen is causing a partial knock-down of the *Mybphl* gene in Cre-positive mice, as opposed to a complete knock-out.

Due to the inverse relationship between MyBP-HL and cMyBP-C [3], both proteins were probed for via immunoblotting **(Figure 1D)**. The MyBP-HL and cMyBP-C signal were compared to total protein, then MyBP-HL was compared to cMyBP-C to calculate a self-normalized ratio. The level of MyBP-HL to total protein in the Cre-positive mice is, on average, half that of the Cre-negative mice. Correspondingly, there is an approximate doubling of the signal for cMyBP-C. The ratios of MyBP-HL:cMyBP-C are significantly lower in the Cre-positive mice, shown in **Figure 1E**. The significantly decreased ratio represents a drop in the MyBP-HL protein levels with a corresponding increase in cMyBP-C levels.

### Conditional Knock-down Mice Have Higher Heart Weights to Body Weights

At the time of tissue collection, measurements for atria, total heart, and body weight were obtained for Cre-negative and Cre-positive Rosa-26-Cre(ERT2) mice ages 11 to 15 wks of age. No significant change is observed in atria to total heart weight or atria to body weight ratios **(Figure 2A)**. Instead, a small yet significant increase in total heart-to-body weight is seen in the Cre-positive mice. When separating the mice out by sex, the higher heart to body weight ratio appears to be primarily due to the male cohort, trending higher with a p-value of 0.0677 by Student’s t-test, as opposed to the female mice which are not significantly different, with a p-value of 0.3789 **(Figure 2B)**. The male cohort is slightly larger than the female, so it is possible that the cohort size disparity is partly responsible for the more pronounced trend in the males. Separating into male and female did not lead to any significance in atrial weight to heart or to body weight comparisons.

**Figure 2.**
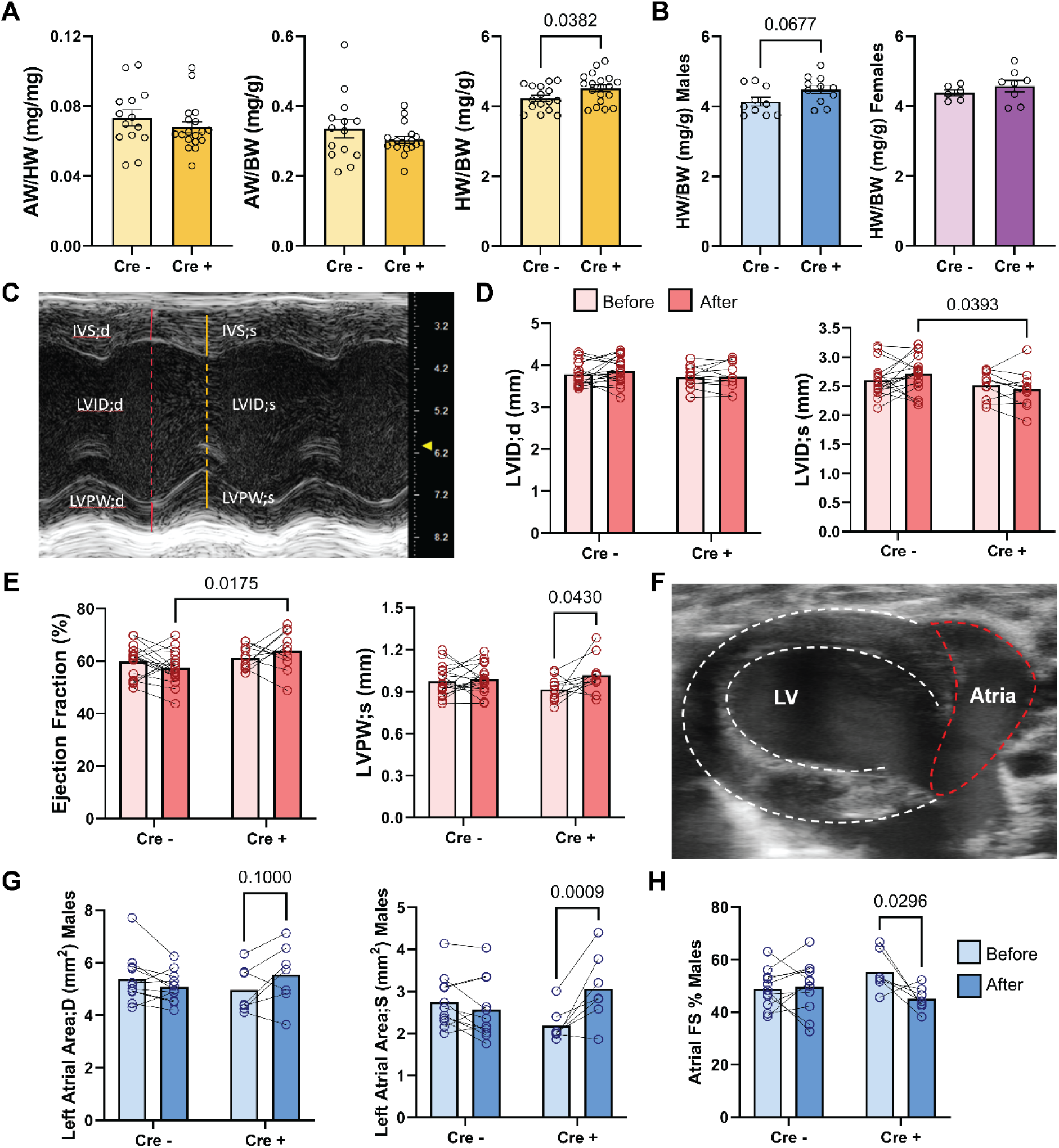
Conditional Reduction of *Mybphl* Causes Cardiac Hypertrophy and Contractile Dysfunction. **A)** Ratios of atrial weight (AW) to total heart weight (HW) or atrial weight to body weight (BW) (Cre-negative: n = 14, Cre-positive: n = 18); total heart weight to body weight is significantly elevated (Cre-negative: n = 16, Cre-positive: n = 19). AW:HW and HW:BW analyzed by Student’s T-test, AW:BW analyzed by Mann-Whitney test after failing Shapiro-Wilks normality test. **B)** Mice were separated into male and female cohorts and re-analyzed, demonstrating a trend toward elevated total heart to body weight ratios in the males (Cre-negative: n = 10, Cre-positive: n = 11) with no significance in the female mice (Cre-negative: n = 6, Cre-positive: n = 8). Data analyzed by Student’s T-test. **C)** Representative M-Mode echocardiography image used to derive left ventricular internal diameter (LVID), interventricular septum diameter (IVS) and Left ventricular posterior wall thickness (LVPW) during peak diastole (;d) and systole (;s). **D)** Quantification of LVID, **E)** ejection fractions and LVPW during systole (;s). Cre (-): n = 18, Cre (+) n = 11. **F)** Representative image of an Electrocardiogram-gated Kilohertz Visualization (EKV) ultrasound, which allows for left atrial area to be measured by tracing atrial border, as shown. **G)** Left atrial area (mm^2^) during diastole and systole in male mice, with the change in atrial area in pre- to post-tamoxifen hearts. **H)** Atrial fractional shortening in pre- and post-tamoxifen measurements. Data analyzed using Student’s t-test (A, B), 2-way ANOVA with repeated measures and Šidák post-hoc analysis, with multiple comparisons performed both within and between groups (D-E, G-H).

### Echocardiography Reveals a Hypercontractile Phenotype

Echocardiograms were performed on mice at 11-12 weeks of age prior to beginning tamoxifen injections, then 6 weeks following. Measurements of left ventricular chamber diameter were taken during peak diastole (relaxation) and peak systole (contraction), shown in **Figure 2C**. Averages of all measurements taken pre- and post-tamoxifen are denoted in **Table 1**. Cre-positive mice overall had a smaller internal diameter of the left ventricle during systole (LVID;s) after tamoxifen treatment than the Cre-negative group, and no change in internal diameter of the left ventricle during diastole (LVID;d) is seen (**Figure 2D)**. Mice in the Cre-positive group additionally have an overall higher ejection fraction and increase in the thickness of the left ventricular posterior wall during systole (LVPW;s) after tamoxifen **(Figure 2E)**. Together, these findings suggest the hearts are becoming hypercontractile. Based on previous findings linking MyBP-HL to DCM [1, 2], the hypercontractile evidence is particularly of interest.

**Table 1:** Conditional Knockdown in Adulthood Promotes a Hypercontractile Phenotype, Primarily in Male Mice. Results from echocardiograms taken directly before and six weeks after tamoxifen injection in Rosa-26-Cre(ERT2) mice. Results were also separated by sex and re-analyzed to look for sex-specific differences. Units are represented as average ± SEM.

|  |  | Cre (-), All mice |  |  |  |  | Cre (+), All mice |  |  |  |  | Cre (-), Males |  |  |  |  | Cre (+), Males |  |  |  |  | Cre (-), Females |  |  |  |  | Cre (+), Females |  |  |  |  |
| --- | --- | --- | --- | --- | --- | --- | --- | --- | --- | --- | --- | --- | --- | --- | --- | --- | --- | --- | --- | --- | --- | --- | --- | --- | --- | --- | --- | --- | --- | --- | --- |
|  | Units | Pre |  | Post |  | n | Pre |  | Post |  | n | Pre |  | Post |  | n | Pre |  | Post |  | n | Pre |  | Post |  | n | Pre |  | Post |  | n |
| HR | bpm | 523 | ±8 | 508 | ±6 | 18 | 505 | ± 10 | 507 | ± 12 | 13 | 524 | ± 12 | 512 | ± 9 | 11 | 500 | ± 14 | 505 | ± 8 | 7 | 521 | ± 11 | 501 | ± 8 | 7 | 512 | ± 15 | 511 | ± 25 | 6 |
| IVS:D | mm | 0.631 | ±0.025 | 0.665 | ±0.027 | 18 | 0.631 | ± 0.022 | 0.603 | ± 0.030 | 11 | 0.639 | ± 0.033 | 0.685 | ± 0.036 | 11 | 0.665 | ± 0.031 | 0.607 | ± 0.038 | 6 | 0.618 | ± 0.042 | 0.634 | ± 0.039 | 7 | 0.590 | ± 0.024 | 0.598 | ± 0.053 | 5 |
| IVS:S | mm | 0.956 | ±0.035 | 0.917 | ±0.035 | 18 | 0.913 | ± 0.025 | 0.946 | ± 0.051 | 11 | 0.972 | ± 0.051 | 0.945 | ± 0.050 | 11 | 0.942 | ± 0.039 | 1.026 | ± 0.065 | 6 | 0.929 | ± 0.044 | 0.873 | ± 0.043 | 7 | 0.879 | ± 0.027 | 0.850 | ± 0.063 | 5 |
| LVID:D | mm | 3.78 | ±0.07 | 3.86 | ±0.08 | 18 | 3.72 | ± 0.08 | 3.73 | ± 0.10 | 11 | 3.90 | ± 0.09 | 4.05 | ± 0.07 | 11 | 3.82 | ± 0.05 | 3.87 | ± 0.10 | 6 | 3.60 | ± 0.06 | 3.57 | ± 0.07 | 7 | 3.60 | ± 0.15 | 3.55 | ± 0.15 | 5 |
| LVID:S | mm | 2.60 | ±0.07 | 2.71 | ±0.07 | 18 | 2.52 | ± 0.07 | 2.45 | ± 0.09 | 11 | 2.68 | ± 0.10 | 2.84 | ± 0.09 | 11 | 2.64 | ± 0.04 | 2.59 | ± 0.11 | 6 | 2.47 | ± 0.07 | 2.51 | ± 0.09 | 7 | 2.37 | ± 0.12 | 2.28 | ± 0.13 | 5 |
| LVPW:D | mm | 0.696 | ±0.026 | 0.721 | ±0.020 | 18 | 0.634 | ± 0.016 | 0.696 | ± 0.033 | 11 | 0.691 | ± 0.029 | 0.738 | ± 0.025 | 11 | 0.648 | ± 0.010 | 0.734 | ± 0.056 | 6 | 0.704 | ± 0.052 | 0.694 | ± 0.035 | 7 | 0.616 | ± 0.033 | 0.650 | ± 0.014 | 5 |
| LVPW:S | mm | 0.976 | ±0.025 | 0.990 | ±0.024 | 18 | 0.915 | ± 0.025 | 1.019 | ± 0.038 | 11 | 0.991 | ± 0.029 | 0.987 | ± 0.031 | 11 | 0.916 | ± 0.033 | 1.054 | ± 0.063 | 6 | 0.952 | ± 0.047 | 0.994 | ± 0.040 | 7 | 0.915 | ± 0.043 | 0.976 | ± 0.035 | 5 |
| EF | % | 59.8 | ±1.5 | 57.5 | ±1.5 | 18 | 61.4 | ± 1.3 | 64.0 | ± 2.2 | 11 | 59.7 | ± 2.0 | 57.4 | ± 2.2 | 11 | 59.2 | ± 1.4 | 62.0 | ± 3.4 | 6 | 59.9 | ± 2.2 | 57.8 | ± 1.9 | 7 | 64.0 | ± 1.7 | 66.3 | ± 2.8 | 5 |
| FS | % | 31.4 | ±1.0 | 29.9 | ±1.0 | 18 | 32.4 | ± 0.9 | 34.4 | ± 1.6 | 11 | 31.4 | ± 1.4 | 30.0 | ± 1.4 | 11 | 30.9 | ± 1.0 | 33.2 | ± 2.4 | 6 | 31.3 | ± 1.5 | 29.8 | ± 1.3 | 7 | 34.1 | ± 1.2 | 35.9 | ± 2.1 | 5 |
| IVRT | ms | 15.9 | ±0.8 | 14.9 | ±0.6 | 18 | 15.5 | ± 0.6 | 14.8 | ± 0.7 | 11 | 13.6 | ± 0.6 | 14.9 | ± 0.8 | 11 | 15.4 | ± 0.8 | 16.5 | ± 2.2 | 7 | 19.4 | ± 0.6 | 15.0 | ± 0.7 | 7 | 17.6 | ± 1.7 | 15.5 | ± 0.9 | 6 |
| E/A | none | 1.52 | ±0.04 | 1.58 | ±0.08 | 18 | 1.52 | ± 0.06 | 1.46 | ± 0.06 | 13 | 1.50 | ± 0.05 | 1.53 | ± 0.08 | 11 | 1.55 | ± 0.09 | 1.48 | ± 0.10 | 7 | 1.55 | ± 0.06 | 1.66 | ± 0.15 | 7 | 1.49 | ± 0.07 | 1.44 | ± 0.06 | 6 |
| E/E' | none | -41.6 | ±6.9 | -42.3 | ±4.5 | 14 | -44.5 | ± 10.4 | -45.2 | ± 4.0 | 11 | -46.6 | ± 8.8 | -47.9 | ± 5.2 | 10 | -61.9 | ± 20.9 | -45.7 | ± 6.8 | 5 | -29.1 | ± 5.4 | -28.5 | ± 3.4 | 4 | -32.2 | ± 3.8 | -43.7 | ± 6.4 | 5 |
| Atrial area:D | mm2 | 5.09 | ±0.22 | 5.07 | ±0.13 | 18 | 4.73 | ± 0.23 | 5.14 | ± 0.34 | 13 | 5.39 | ± 0.28 | 5.09 | ± 0.18 | 11 | 4.96 | ± 0.33 | 5.54 | ± 0.44 | 7 | 4.62 | ± 0.29 | 5.05 | ± 0.20 | 7 | 4.45 | ± 0.31 | 4.67 | ± 0.48 | 6 |
| Atrial area:S | mm2 | 2.65 | ±0.12 | 2.52 | ±0.14 | 18 | 2.21 | ± 0.11 | 2.72 | ± 0.22 | 13 | 2.75 | ± 0.19 | 2.57 | ± 0.22 | 11 | 2.19 | ± 0.15 | 3.07 | ± 0.31 | 7 | 2.50 | ± 0.11 | 2.43 | ± 0.13 | 7 | 2.23 | ± 0.17 | 2.30 | ± 0.21 | 6 |
| Atrial FS% | % | 47.5 | ±1.5 | 50.3 | ±2.2 | 18 | 52.5 | ± 2.5 | 47.2 | ± 2.0 | 13 | 48.9 | ± 2.2 | 49.8 | ± 3.0 | 11 | 55.3 | ± 2.8 | 45.1 | ± 1.7 | 7 | 45.3 | ± 1.5 | 51.2 | ± 3.6 | 7 | 49.1 | ± 4.0 | 49.6 | ± 3.8 | 6 |

### Atrial Contractility Defects are Noted on Echocardiography

In tandem with the ventricular deficits seen, the Cre-positive mice have atrial dysfunction. Left atrial areas were measured using tracings similar to that shown in **Figure 2F**. After separating into male vs female, it was noted that all significant findings are driven by the male cohort, whereas the female mice do not demonstrate these atrial changes. Cre-positive male mice have a significant increase in left atrial area during systole after tamoxifen compared to before. The left atrial area during diastole also seems to trend upward **(Figure 2G)**. Atrial fractional shortening was calculated to determine how well the left atria are contracting using the following equation:

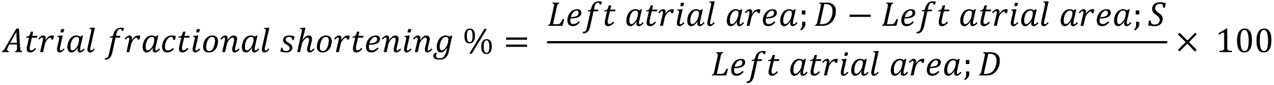

The Cre-positive male mice have a statistically significant decrease in atrial fractional shortening compared to the Cre-negative mice **(Figure 2H)**. This finding is in keeping with the increase in left atrial area during systole and indicates that Cre-positive male mice have hypocontractile left atria that may additionally be trending toward enlargement during diastole.

### Conditional Knock-down Causes a Worse Phenotype in Male Mice than Female Mice

The changes to LVPW;s noted in **Figure 2** appear to largely be associated with changes in the male mice, as this value trends toward significance when comparing just males and does not approach significance in the females. The LVPW;s is in males has a p-value of 0.059 for Cre-positive Before vs After, and in females the p-value is 0.599. This pattern carries over to evaluation of atrial function, wherein the comparison of left atrial area during diastole and systole, and thus atrial fractional shortening, all demonstrate statistically significant changes in the male mice with no changes seen between female cohorts **(Table 1)**. Interestingly, female Cre-positive mice have a significantly higher fractional shortening compared to female Cre-negative mice, and male mice have no significant changes in this metric.

### Telemetry Monitoring Demonstrates Increased P-wave Duration in Cre-positive Mice

Mice were observed under baseline conditions, then after treatment with isoproterenol, propranolol, and flecainide on separate days. The durations of the P-waves are significantly elevated after isoproterenol and propranolol treatment for all Cre-positive individuals and trend upward after flecainide treatment (**Figure 3D**). Additionally, a trend toward decrease in the QRS complex duration is observed after isoproterenol and flecainide **(Figure 3E)**. No changes in RR interval duration were noted between Cre-negative and Cre-positive mice within the various drug treatments. A trend of RR interval increase was seen under propranolol conditions; however, the Cre-positive sample size is three mice (n = 3) and this was largely drawn upward by one particular Cre-positive mouse that had an incredibly varied heart rate. A rhythm strip can be seen corresponding to this mouse in **Figure 3A**. Although unclear exactly what rhythm this mouse is experiencing, it appears to potentially be a sick-sinus syndrome or an intermittent junctional rhythm. In panel **3B**, the morphology of the P-waves can be seen changing from beat to beat and at times a P-wave is difficult to identify at all, suggesting a potential ectopic pacemaker/junctional beat. The RR-interval oscillates in waves, as seen in the bottom part of panel **3A**. The RR interval for one beat to the next corresponding beat was plotted as a Poincaré plot with a representative Cre-negative mouse also shown for comparison (**Figure 3C),** demonstrating the high amount of RR interval variability in this Cre-positive mouse that is not seen in control mice.

**Figure 3.**
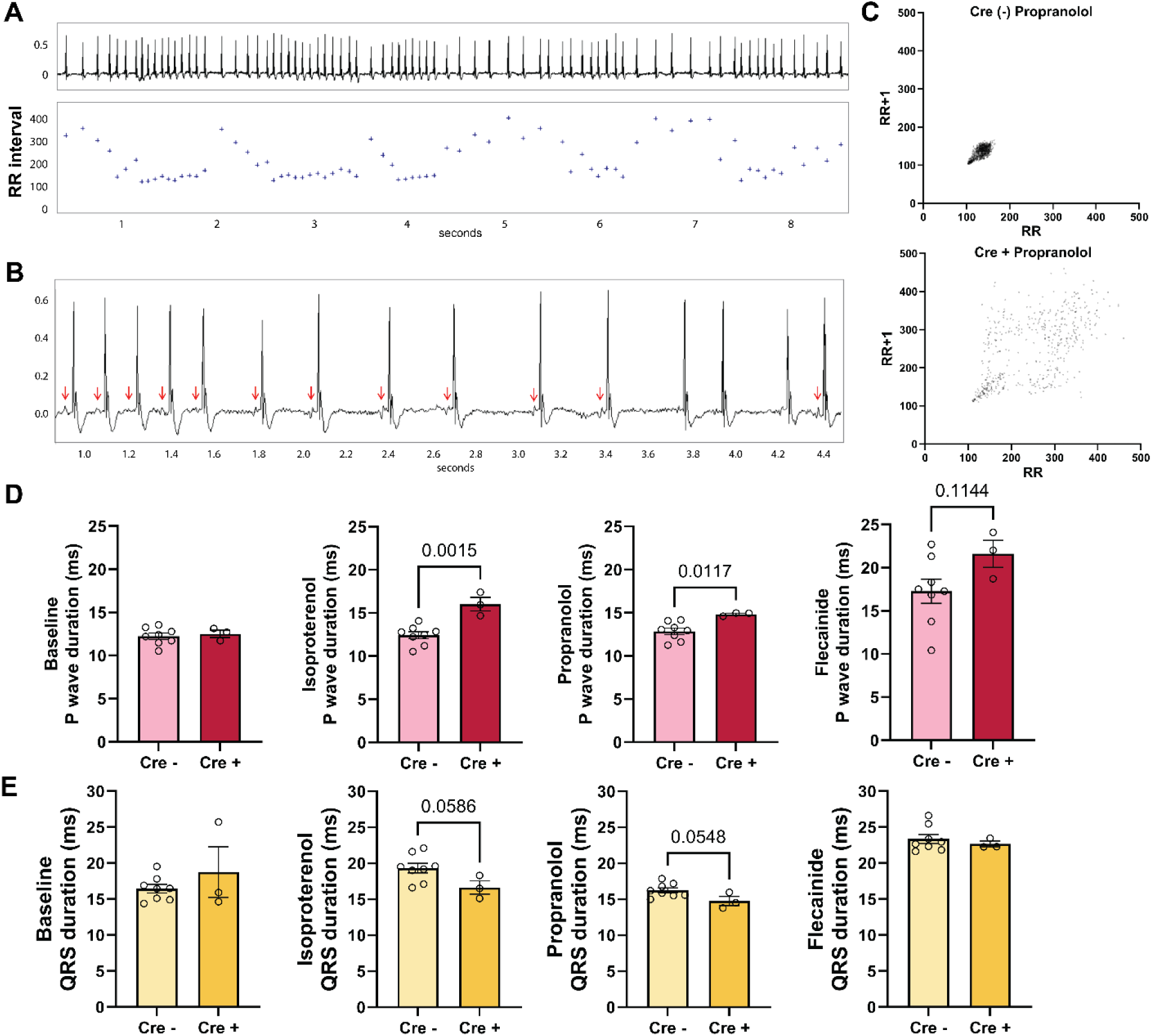
Conditional Knock-down of MyBP-HL in Adulthood Causes Conduction Abnormalities, Including Increased P-wave Duration. A) A portion of the ECG from a Cre-positive mouse demonstrates significant variability in heart rate. The RR interval, shown underneath the strip in milliseconds, appears to oscillate in frequency. B) A close-up view of the mouse’s rhythm demonstrates alterations in p-wave morphology during moments when the beats have abnormally long RR intervals. C) Poincaré plots demonstrate the variability from one RR interval to the one directly following it. The representative plots shown indicate an extremely high RR variability within the Cre-positive mouse after propranolol administration. Each plot contains 475 data points. D-E) Cre-negative (n = 8) and Cre-positive (n = 3) mice were observed under the following conditions: baseline (overnight), isoproterenol, propranolol, and flecainide. In both isoproterenol and propranolol treatments, the Cre-positive cohort has a significantly longer P-wave duration and trend toward shorter QRS complex duration. Data were analyzed by Students’ t-tests.

### Confirming conduction system specific loss of Mybphl

DNA isolated from the right atria of Cntn-2-Cre mice was amplified using forward primer 1 and the reverse primer marked in **Figure 1B**. Cre-positive mice demonstrate a band at 270 bp that Cre-negative mice do not have, representing the recombined *Mybphl* gene **(Figure 4A)**.

**Figure 4.**
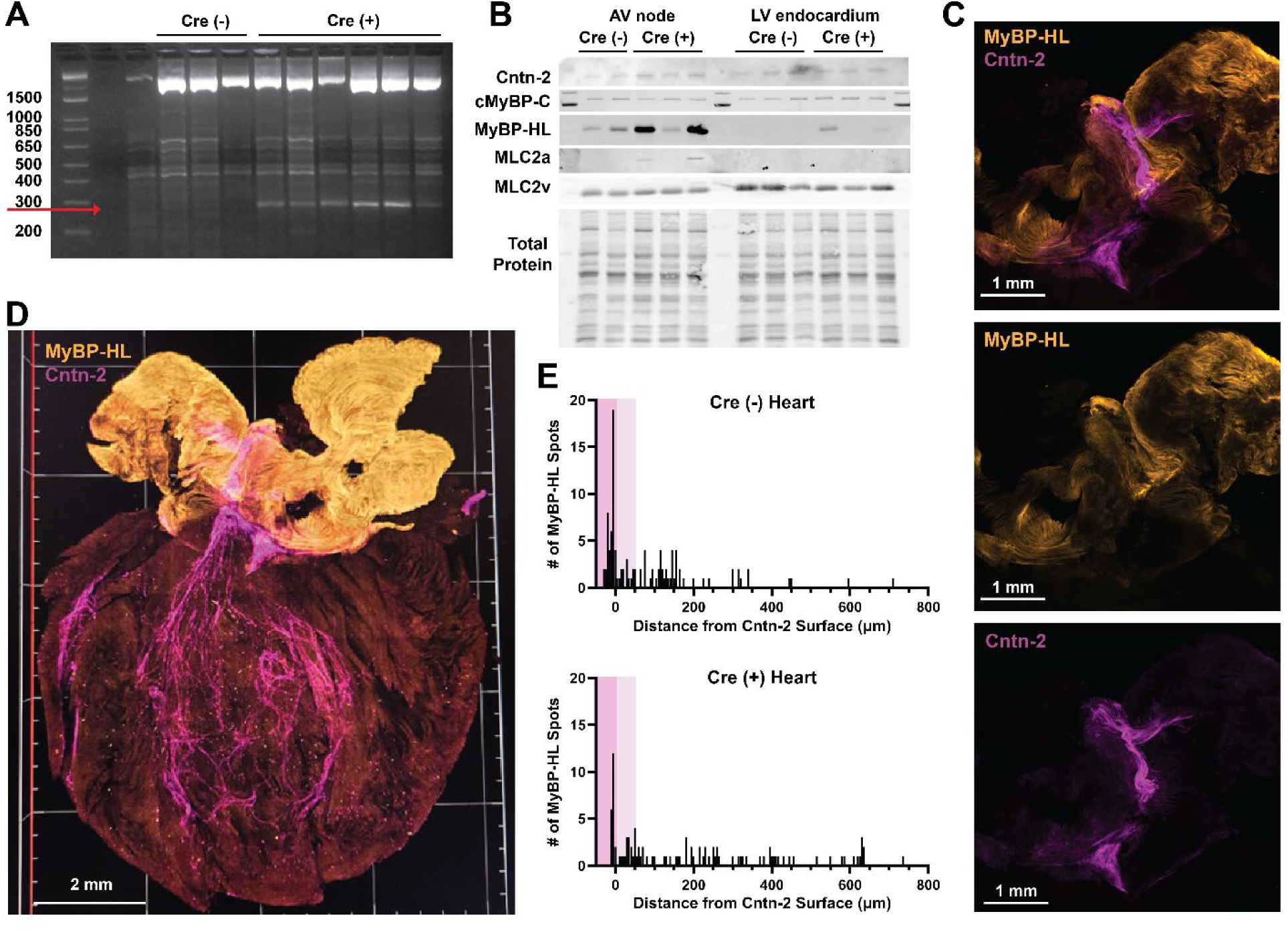
Contactin-2-Cre Mice Cause Conditional Knockout of *Mybphl* in the Conduction System. A) PCR gel to amplify the recombined *Mybphl* gene, showing a band in the Cre (+) samples (n = 6) and not in the Cre (-) samples (n = 3). B) Immunoblot with AV node and left ventricular (LV) endocardium tissue was performed. The blot was incubated with antibodies for Cntn2, cMyBP-C, MyBP-HL, myosin light chain 2, atrial isoform (MLC2a), and myosin light chain 2, ventricular isoform (MLC2v). For AV node samples, Cre (-) n = 2, Cre (+) n = 3. For LV endocardium, n = 3 for both. C-D) Mouse heart tissue was cleared and incubated with an antibody for MyBP-HL (orange) and Contactin-2 (purple). A 3D rendering was constructed based on acquired signal. **C)** The atria of the Cntn-2-Cre positive mouse (shown) depicts the path the atrial conduction system carves through the atria before reaching the AV node. MyBP-HL, shown in orange, has areas of decreased signal seen within the atria. These regions correspond to the areas with high Contactin-2 signal (purple). D) A mouse heart was cut coronally and tissue cleared. In the ventricles, the MyBP-HL puncta can be visualized. E) The distance in μm from each MyBP-HL spot to the Cntn-2 surface was measured and plotted in a histogram. Each bin is 5 μm. The Cre-negative heart, which did not undergo recombination, has more MyBP-HL spots that are less than zero μm away from the nearest Cntn-2 surface (darker pink highlight), indicating colocalization. No overall change was noted in the number of spots directly next to the Cntn-2-Cre surface (light pink), demonstrating the specificity of the knock-out within the model.

Due to the extremely low portion of MyBP-HL cells being deleted, it proved difficult to analyze level of protein recombination. Tissue from the AV node region and left ventricular (LV) endocardium were analyzed using immunoblotting, as shown in **Figure 4B**. The higher levels of MyBP-HL in the AV node tissue are attributed to the signal increase in myosin light chain 2a (MLC2a), the atrial isoform, indicating some atrial tissue was collected in the sample. Since *Mybphl* is expressed at 10,000 times higher levels in the atria [1], any inclusion of atrial tissue will largely affect MyBP-HL signal. The amount of cMyBP-C does trend upward in the Cre-positive samples of the LV endocardium, but not significantly. MyBP-HL signal is also slightly higher in these samples, however presence or absence of MyBP-HL in and of itself is not a great indicator of recombination. Some MyBP-HL foci in the ventricular are within Contactin-2 expressing cells, but also near or adjacent to the cells [2]. Therefore, if the portion of tissue selected for homogenization had more foci that are not located in Contactin-2 expressing cells, MyBP-HL expression may be higher on immunoblotting. The apparent heterogeneity and seemingly random pattern of expression is another confounding variable involved in the excision of tissue regions for sample preparation.

### Airyscan Imaging Demonstrates a Reduction in MyBP-HL Colocalization with Contactin-2

One Contactin-2-Cre negative and one Cre-positive heart underwent tissue clearing after being cut in half along the coronal plane. Samples were incubated with antibodies specific for MyBP-HL and Contactin-2, as a marker for the conduction system. These were imaged to create a 3D rendering of half of each of these hearts. The rendering created for the Contactin-2-Cre positive heart can be seen in **Figure 4D**.

The Imaris software was used to pick out MyBP-HL positive spots within the ventricle, and the distance from these spots to the Purkinje fiber surface was calculated. The distributions of MyBP-HL spots within these two hearts compared to a surface rendering of Cntn-2 signal are plotted in **Figure 4E**. In the Cre-negative heart, 112 MyBP-HL expressing puncta are identified, 45 of which are within the labeled Purkinje fiber region. In the Cre-positive heart, of the 101 spots identified, only 19 are localized within the Contactin-2 labeled region.

Seeing as Cntn-2 is also expressed within the atrial conduction pathways, qualitative views of a subregion of the atria were obtained. When the Cre-positive heart is separated into its respective channels in **Figure 4C**, MyBP-HL signal, seen in orange, appears fainter in the regions with high Cntn-2 signal (purple). Altogether, the evidence demonstrates that although MyBP-HL foci still exist in the ventricle in Cntn-2-Cre positive mice, the amount directly within the conduction system is substantially decreased.

### Cntn-2-Cre Mice Show No Significant Changes in Heart Mass

The Cntn-2-Cre positive mice were weighed at time of tissue collection in a similar manner to the Rosa26-Cre(ERT2) mice. Both heterozygous, containing one wild type allele and one floxed MyBP-HL allele (WT/Fl), and homozygous floxed/floxed (Fl/Fl) mice are used in experiments. Cre-negative mice from both WT/Fl and Fl/Fl backgrounds are represented in the Cre-negative column. Across the different groups, no significant differences are noted in atrial weight to total heart weight, atrial weight to body weight, or heart weight to body weight **(Figure 5A)**. The atrial weights for the Fl/Fl + cohorts have several values that are higher than the rest of the samples. The Fl/Fl + group did not pass a normality check for a Gaussian distribution for the Atrial Weight to Total Heart Weight and Atrial Weight to Body Weight analyses. Thus, a Kruskal-Wallis test was performed for these two comparisons. The cohorts were also separated by sex, and no significant changes are seen within sex-specific cohorts.

**Figure 5.**
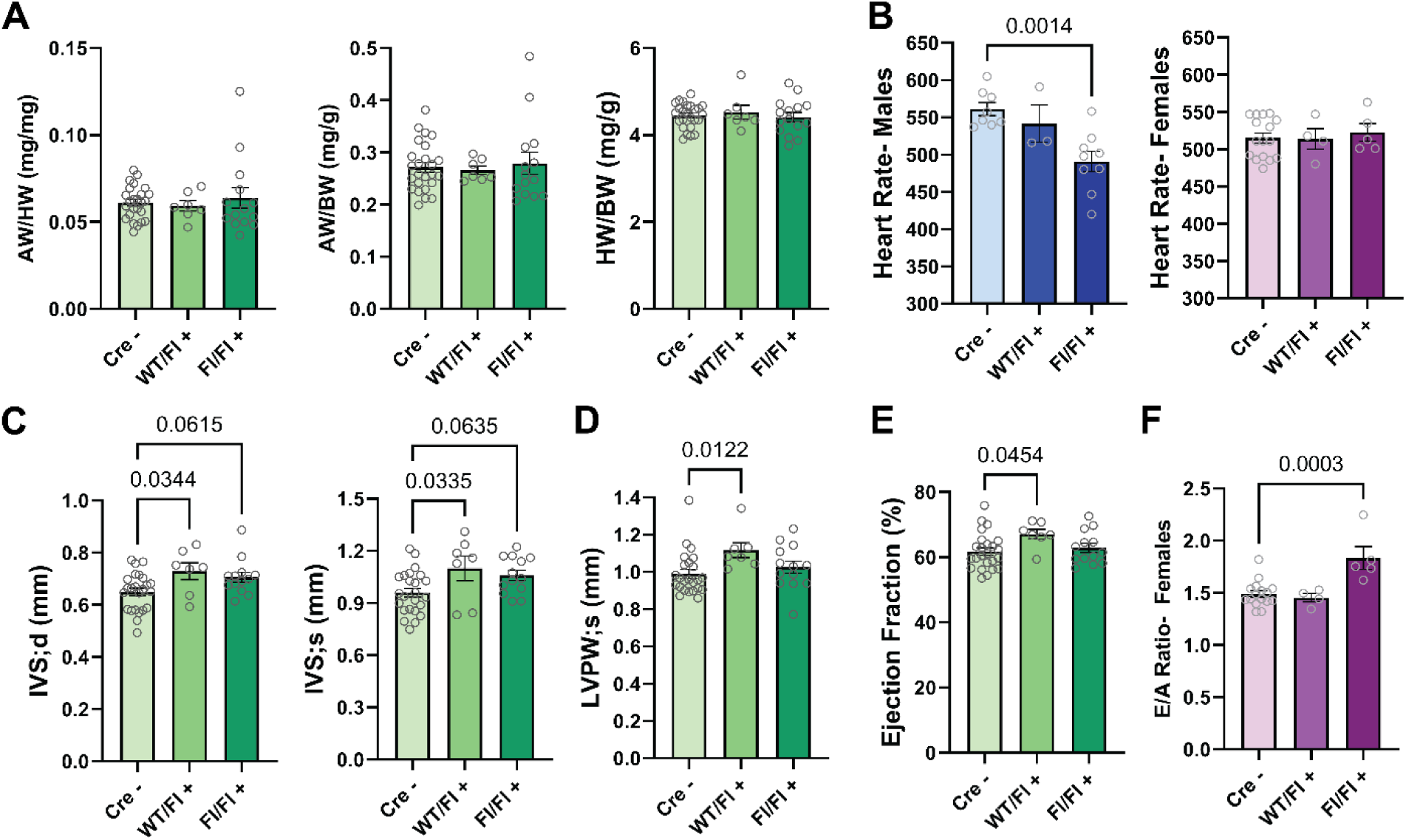
Heart Mass and Ventricular Function Are Not Altered After MyBP-HL Deletion in the Conduction System. A) Across the various cohorts, atrial weight (AW) is compared to total heart weight (HW) and body weight (BW) (Cre (-) n = 25, WT/Fl (+) n = 7, Fl/Fl (+) n = 14), and total heart weight is also analyzed against body weight (Cre (-) n = 25, WT/Fl (+) n = 7, Fl/Fl (+) n = 15). No significant differences are noted between the cohorts. Data are analyzed by Kruskal-Wallis test for AW/HW and AW/BW, and by one-way ANOVA for HW/BW. B) Heart rate is significantly decreased in the male Fl/Fl Cre (+) cohort compared to the male Cre (-) group (Cre (-) n = 8, WT/Fl (+) n = 3, Fl/Fl (+) n = 9), but this difference is not seen in the female mice (Cre (-) n = 16, WT/Fl (+) n = 4, Fl/Fl (+) n = 5). C) The IVS thickness during diastole and systole is significantly increased in the WT/Fl cohort and trending upward in the Fl/Fl cohort. D-E) The WT/Fl cohort shows an increased thickness of the left ventricular posterior wall with increased ejection fraction percent. Data are analyzed by one-way ANOVA with multiple comparison using the Cre (-) group as the control. LVPW;s is analyzed by Kruskal-Wallis. Cre (-) n = 24, WT/Fl (+) n = 7, and Fl/Fl (+) n = 13.

### Cardiac Conduction System Deletion Induces Decreased Heart Rate and an Increase in Thickness of the Ventricular Walls

Echocardiography was performed on the Contactin-2-Cre mice around 18 weeks of age, approximately the same age as the after-tamoxifen echocardiograms on the Rosa-26-Cre(ERT2) mice. The male Fl/Fl Cre-positive mice had lower average heart rates during image acquisition under isoflurane compared to the male Cre-negative mice **(Figure 5B)**. On average, their heart rates were 50 bpm lower than their Cre-negative counterparts, seen in **Table 2**.

**Table 2:** Conditional Knock-out Within Cardiac Conduction System Leads to Decreased Heart Rate and Thickened Ventricular Walls. Results from echocardiograms taken at 18 weeks of age in Contactin-2-Cre mice. Results were separated by sex and re-analyzed to look for sex-specific differences. Units are represented as average ± SEM.

|  |  | All Cre (-) |  |  | All WT/FI Cre (+) |  |  | All FI/FI Cre(+) |  |  | Male Cre (-) |  |  | Male WT/FI Cre (+) |  |  | Male FI/FI Cre(+) |  |  | Female Cre (-) |  |  | Female WT/FI Cre (+) |  |  | Female FI/FI Cre(+) |  |  |
| --- | --- | --- | --- | --- | --- | --- | --- | --- | --- | --- | --- | --- | --- | --- | --- | --- | --- | --- | --- | --- | --- | --- | --- | --- | --- | --- | --- | --- |
|  | Units | Ave | SEM | n | Ave | SEM | n | Ave | SEM | n | Ave | SEM | n | Ave | SEM | n | Ave | SEM | n | Ave | SEM | n | Ave | SEM | n | Ave | SEM | n |
| HR | bpm | 530 | ±7 | 24 | 526 | ±13 | 7 | 502 | ±10 | 14 | 561 | ±9 | 8 | 542 | ±25 | 3 | 491 | ±14 | 9 | 515 | ±7 | 16 | 514 | ±14 | 4 | 523 | ±12 | 5 |
| IVS;D | mm | 0.650 | ±0.015 | 24 | 0.728 | ±0.032 | 7 | 0.706 | ±0.019 | 13 | 0.708 | ±0.015 | 8 | 0.767 | ±0.020 | 3 | 0.733 | ±0.026 | 8 | 0.621 | ±0.017 | 16 | 0.699 | ±0.053 | 4 | 0.664 | ±0.017 | 5 |
| IVS;S | mm | 0.96 | ±0.03 | 24 | 1.10 | ±0.07 | 7 | 1.06 | ±0.03 | 13 | 1.07 | ±0.04 | 8 | 1.19 | ±0.02 | 3 | 1.11 | ±0.03 | 8 | 0.91 | ±0.02 | 16 | 1.03 | ±0.12 | 4 | 0.98 | ±0.03 | 5 |
| LVID;D | mm | 3.72 | ±0.05 | 24 | 3.72 | ±0.12 | 7 | 3.76 | ±0.10 | 13 | 3.86 | ±0.05 | 8 | 3.93 | ±0.18 | 3 | 3.95 | ±0.12 | 8 | 3.64 | ±0.06 | 16 | 3.55 | ±0.11 | 4 | 3.46 | ±0.07 | 5 |
| LVID;S | mm | 2.50 | ±0.05 | 24 | 2.36 | ±0.09 | 7 | 2.50 | ±0.07 | 13 | 2.51 | ±0.08 | 8 | 2.44 | ±0.14 | 3 | 2.62 | ±0.06 | 8 | 2.49 | ±0.06 | 16 | 2.30 | ±0.12 | 4 | 2.30 | ±0.10 | 5 |
| LVPW;D | mm | 0.699 | ±0.021 | 24 | 0.727 | ±0.040 | 7 | 0.709 | ±0.026 | 13 | 0.742 | ±0.058 | 8 | 0.786 | ±0.085 | 3 | 0.708 | ±0.023 | 8 | 0.677 | ±0.013 | 16 | 0.683 | ±0.021 | 4 | 0.711 | ±0.062 | 5 |
| LVPW;S | mm | 0.99 | ±0.02 | 24 | 1.12 | ±0.04 | 7 | 1.03 | ±0.03 | 13 | 1.08 | ±0.05 | 8 | 1.19 | ±0.08 | 3 | 1.04 | ±0.04 | 8 | 0.95 | ±0.02 | 16 | 1.07 | ±0.02 | 4 | 1.00 | ±0.07 | 5 |
| EF | % | 61.8 | ±1.1 | 24 | 67.0 | ±1.5 | 7 | 62.8 | ±1.4 | 13 | 64.9 | ±2.3 | 8 | 68.7 | ±1.2 | 3 | 62.7 | ±1.3 | 8 | 60.3 | ±1.1 | 16 | 65.8 | ±2.4 | 4 | 63.0 | ±3.2 | 5 |
| FS | % | 32.7 | ±0.8 | 24 | 36.5 | ±1.1 | 7 | 33.5 | ±1.0 | 13 | 35.1 | ±1.7 | 8 | 37.9 | ±0.9 | 3 | 33.4 | ±1.0 | 8 | 31.5 | ±0.8 | 16 | 35.5 | ±1.7 | 4 | 33.5 | ±2.4 | 5 |
| IVRT | ms | 15.3 | ±0.7 | 24 | 13.1 | ±1.3 | 7 | 14.7 | ±0.9 | 14 | 12.6 | ±0.7 | 8 | 12.0 | ±2.4 | 3 | 13.5 | ±1.1 | 9 | 16.6 | ±0.7 | 16 | 13.8 | ±1.5 | 4 | 16.9 | ±0.7 | 5 |
| E/A | none | 1.47 | ±0.03 | 24 | 1.40 | ±0.04 | 7 | 1.56 | ±0.08 | 14 | 1.44 | ±0.08 | 8 | 1.33 | ±0.04 | 3 | 1.40 | ±0.06 | 9 | 1.49 | ±0.03 | 16 | 1.45 | ±0.04 | 4 | 1.83 | ±0.11 | 5 |
| E/E' | none | -44.4 | ±4.9 | 23 | -41.5 | ±6.8 | 6 | -53.6 | ±7.0 | 12 | -66.1 | ±12.0 | 7 | -79.0 | ±19.6 | 2 | -45.6 | ±6.6 | 8 | -34.9 | ±2.5 | 16 | -40.8 | ±4.5 | 4 | -33.4 | ±3.6 | 4 |
| Atrial area;D | mm<sup>2</sup> | 4.99 | ±0.19 | 24 | 4.66 | ±0.18 | 7 | 5.06 | ±0.20 | 13 | 5.90 | ±0.28 | 8 | 5.05 | ±0.18 | 3 | 5.33 | ±0.22 | 9 | 4.53 | ±0.17 | 16 | 4.37 | ±0.17 | 4 | 4.46 | ±0.21 | 4 |
| Atrial area;S | mm<sup>2</sup> | 2.51 | ±0.12 | 24 | 2.45 | ±0.17 | 7 | 2.41 | ±0.14 | 13 | 2.99 | ±0.21 | 8 | 2.45 | ±0.45 | 3 | 2.56 | ±0.17 | 9 | 2.27 | ±0.10 | 16 | 2.46 | ±0.06 | 4 | 2.08 | ±0.22 | 4 |
| Atrial FS% | % | 49.5 | ±1.5 | 24 | 47.1 | ±3.6 | 7 | 52.4 | ±2.0 | 13 | 49.5 | ±2.2 | 8 | 51.9 | ±8.0 | 3 | 52.0 | ±2.1 | 9 | 49.6 | ±1.9 | 16 | 43.5 | ±2.1 | 4 | 53.3 | ±5.0 | 4 |

The thickness of the interventricular septum (IVS) is significantly higher in the WT/Fl group and trending higher in the Fl/Fl mice, during both diastole and systole **(Figure 5C)**. The left ventricular posterior wall thickness during systole (LVPW;s) is higher in the WT/Fl group but not in the Fl/Fl mice **(Figure 5D)**. The data distributions for LVPW;s during systole does not follow a normal distribution, per Shapiro-Wilk testing, and thus this data set was analyzed by Kruskal-Wallis. The WT/Fl group also has significantly increased ejection fraction compared to Cre-negative mice **(Figure 5E)**. Separating into male and female does not demonstrate a clear sex-specific trend for these parameters. The E/A ratio, which was not significant in a mixed-sex analysis, was significantly increased in the female Fl/Fl mice compared to the Cre-negative female mice, with a p-value of 0.0003 **(Figure 5F)**. Taken together, we see a similar trend as the conditional knock-down mice, wherein the loss of MyBP-HL in the conduction system pushes these hearts toward a hypercontractile phenotype.

### Telemetry Analysis Does Not Show Any Overt Changes in Cardiac Rhythm

Between the Contactin-2-Cre negative and positive mice, no significant changes in average QRS interval, PR interval, P wave duration, or RR interval are noted **(Figure 6B-C)**. Based on the Poincaré plots generated, it does seem that Cre-positive mice have slightly higher variability in their RR intervals after isoproterenol treatment and propranolol treatment **(Figure 6A)**. In total, 4 Cre-negative and 4 Cre-positive mice were analyzed. Each cohort had 2 male mice and 2 female mice in it. The data was separated by sex and re-analyzed with no statistically significant findings. The group size is relatively small, however, when separated by sex.

**Figure 6.**
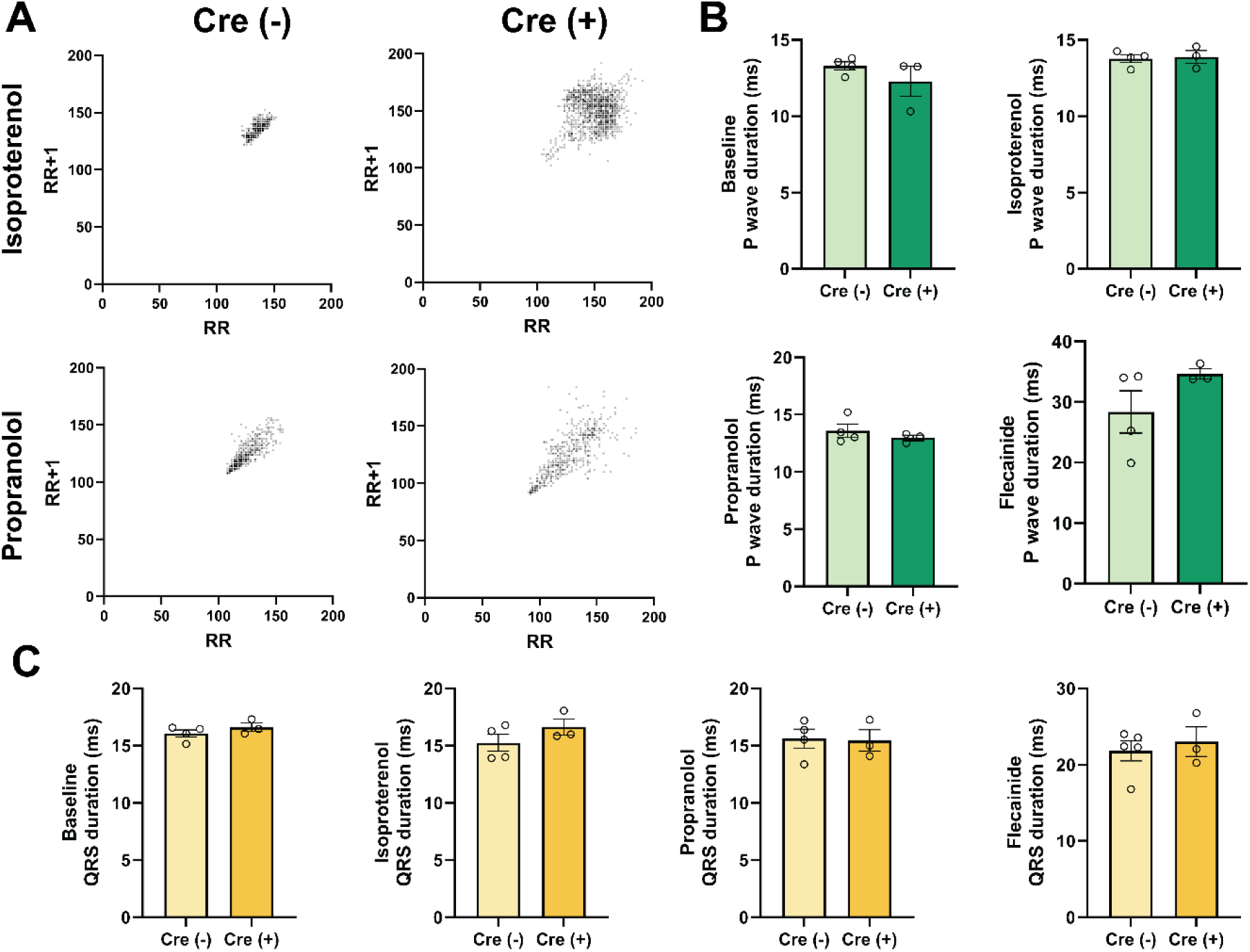
Loss of MyBP-HL in the Conduction System Does Not Affect P-wave Duration, But Seems to Increase Beat-to-Beat Variability. A) Representative Poincaré plots of Cre negative (left) and Fl/Fl Cre-positive (right) mice after treatment with isoproterenol and propranolol. The Cre-positive mice appear to have a wider spread of beat variabilities compared to Cre-negative mice. Each plot contains 500 data points. B-C) Cre-negative (n = 4) and Cre-positive (n = 3) mice were observed under the following conditions: baseline (overnight), isoproterenol, propranolol, and flecainide. No significant differences were noted between the two cohorts for P wave duration or QRS complex duration.

## DISCUSSION

Myosin Binding Protein H-Like is a recently discovered sarcomere protein expressed in the heart. MyBP-HL loss of function mutations in humans are associated with a DCM phenotype, with thinned ventricular walls and poor contractility in the ventricles. Moreover, atrial dilation and an increased propensity for a multitude of arrhythmias are seen [1]. The causes and triggers of DCM are still poorly understood; therefore, characterization of this protein and understanding the mechanism by which MyBP-HL mutations cause disease could lead to a better understanding of DCM as a whole. This knowledge may provide opportunities to develop better therapies for treating the underlying cause of DCM, instead of just managing symptoms [9].

### Conditional Knock-down of Mybphl in Adult Mice Causes a Hypertrophic Phenotype

A previously characterized mouse model with constitutive deletion of MyBP-HL recapitulates the phenotype seen in humans, in both heterozygous and homozygous null mouse models [1]. We analyze whether MyBP-HL is necessary for proper cardiac function in a model that decreases the level of MyBP-HL in adulthood. If knock-down were to not cause disease in adult mice, this could implicate the protein for a role in cardiac development.

The Rosa26-Cre(ERT2) model creates a partial knock-down of the MyBP-HL protein after tamoxifen administration. We chose the time frame of around 11 to 12 weeks to start tamoxifen injections, when the mice are fully grown. Immunoblotting confirmed the model not only a has decreased MyBP-HL protein levels, but also a corresponding increase in the level of cMyBP-C, which shares an inverse relationship with MyBP-HL and competes for binding sites along the sarcomere thick filament [3].

Our results demonstrate that conditional knock-down of MyBP-HL in adult mice is correlated with a hypertrophic phenotype. The Cre-positive mice demonstrate increased fractional shortening and ejection fraction, a larger left ventricular internal diameter during systole, and thickening of the left ventricular posterior wall during systole, as summarized in **Table 1**, indicating a hypercontractile phenotype. The conditional knock-down also correlates with increased total heart to body weight ratio, as would be expected in a hypertrophic heart. This data contrasts with the DCM phenotype observed in the constitutive knock-out model [1]. Similar to the prior constitutive model is the impaired atrial contraction, seen by the increased left atrial area during diastole and systole and the lower atrial fractional shortening, particularly in male mice [1].

When the cohorts are separated by sex and re-analyzed, clear sex-specific differences are seen. In terms of physical changes, male mice trend toward significance for a decrease in left ventricular internal diameter and increased left ventricular posterior wall thickness during systole and show a significant increase in left atrial area during diastole and systole. Female mice do not show any significant changes or even a trend toward significance in any of these physical parameters. Yet, when looking at functional changes, the female mice have a slightly more significant increase in fractional shortening than males. This suggests that the structural remodeling may impact male mice first or more severely than female mice.

It is known that in humans, DCM commonly affects more males than females at an estimated 3:1 ratio [10, 11]. This does however indicate that the diagnosis, prognosis, and treatment criteria for DCM are all informed by DCM presentation in males, more so than it is in females. Recently, it was found that in female DCM patients with less dilation of their ventricles and higher ejection fraction were more likely to experience a major heart failure event, up to and including cardiovascular death [12]. The statistics for hypertrophic cardiomyopathy (HCM) are similar in terms of prevalence, with approximately 60% of all HCM patients being male [13]. The average age of diagnosis in female patients is also older than that of male patients, however female patients with HCM usually have a worsened phenotype and prognosis by the time they are diagnosed. The age of diagnosis does appear to have a genetic component. Patients with an *MYH7* mutation had no difference in age of diagnosis between males and females, however female patients with *MYBPC3* mutations and thin-filament mutations were on average 4.8 and 6.7 years older, respectively, than males with the same mutations [13]. This proposes an interesting link to sex differences for MyBP-HL as well due to its high homology with cMyBP-C and provides a potential use for this model in determining the specific pathway that may be conferring cardiac protection to female mice. On the other hand, it may be that the current parameters used to diagnose HCM and DCM, which are largely based on overt physical changes, are not the best gauges of disease in females, and more work needs to be done to explore other potential diagnostic criteria, with the potential for this model to be of use in this investigation.

### Conditional Knock-down of Mybphl in Adult Mice Is Sufficient to Induce Conduction Changes

An increased P-wave duration is observed in the Cre-positive mice with both isoproterenol and propranolol administration that is not present at baseline, with a trend upward present after flecainide administration. This increased duration indicates that the time for the depolarization signal to spread through the atria is slower with partial loss of MyBP-HL, providing evidence that this contractile protein is essential for proper conduction of electrical impulses through atrial cardiomyocytes. The constitutive knock-out models previously characterized had a significant increase in the QRS duration with isoproterenol, demonstrating a similar slowing of electrical signal, but in the ventricle as opposed to the atria [1]. Interestingly, the conditional knock-down demonstrates a trend toward a decrease in QRS duration after isoproterenol and propranolol, the exact opposite of the constitutive knock-out. Both isoproterenol and propranolol work on beta receptors in the heart and are related to adrenergic signaling. Whereas isoproterenol is a beta receptor agonist, propranolol blocks beta receptors [14, 15]. Flecainide, which acts on sodium channels in the heart [16], had a lesser effect on electrical changes, providing evidence that the link between MyBP-HL and cardiac conduction relates to adrenergic signaling.

The Cre-positive mice that underwent conditional knock-down also appeared to have increased RR variability, with one of the three mice in particular demonstrating wide oscillations in its RR intervals. Healthy mice do not experience this level of RR variability under normal conditions, indicating this Cre-positive mouse has a conduction abnormality most likely due to the decrease in MyBP-HL. This is further supported by the variation in P-wave morphology on the close-up view of the rhythm strip, showing that the arrhythmia appears to be originating in the atria.

### Cardiac Conduction System Knock-Out Causes Lower Heart Rate and Ventricular Wall Hypertrophy

The expression pattern of MyBP-HL is uncommon in that it exists in specific puncta within the ventricles, congregating around the ventricular conduction system [1, 2]. MyBP-HL is shown to be enriched within a subset of cells related to the transition between the conduction fibers and the myocardium [17]. The need for differential thick filament regulation within this subset of cells is unclear, and we explored what the consequences of deleting MyBP-HL solely in the cardiac conduction system would be for cardiac function.

Ventricular arrhythmias are common in DCM, yet the gene responsible for causing this phenotype is typically associated with cellular processes unrelated to sarcomeric contraction. In patients with DCM, 33% of all cases of atrioventricular (AV) block are caused by mutations in Lamin A/C, despite it making up only 6% of all DCM cases [18]. While many arrhythmias in DCM can be linked to remodeling in the heart [18], the proband in which MyBP-HL was discovered and two of his siblings had congenital third-degree AV block in childhood, prior to the onset of DCM [1]. Therefore, it appears that MyBP-HL has an effect on the conduction system separate from its ability to influence the development of whole-heart contractile deficits.

The Contactin-2-Cre mouse deletes *Mybphl* solely in the cardiac conduction system of the heart to allow for evaluation of the role of the MyBP-HL protein in conduction. One noted finding is that the Fl/Fl Cre-positive mice had lower heart rates during their echocardiograms. The telemetry analysis did not reveal any differences in heart rate. The lower rates during the echoes may have been related to an effect of the isoflurane on the Cre-positive hearts that is not seen at baseline, or it could be the case that the sample size of four Cre-negative and three Cre-positive mice for the telemeters was underpowered. The decreased heart rate was almost exclusively seen in the male mice, and the telemeters were implanted in a mixed cohort, but these telemetry studies were not powered to identify these changes. Delayed conduction through the atria was noted in the Rosa26-Cre(ERT2) mice, and the constitutive knock-out model had prolonged QRS complexes[1], so a lower heart rate when MyBP-HL is deleted from the conduction system is consistent with other evidence. If this is caused by a slow conduction pathway, it could explain the higher propensity for arrhythmias or any ectopic pacemakers. If it is taking too long for a signal to reach any atrial or ventricular foci, they may start trying to pace the heart themselves, causing ectopy. The higher QRS interval in the constitutive null model is potentially caused by the increased distance the signal needs to travel through in a dilated heart. This finding supports the hypothesis that the puncta of MyBP-HL expressing cells associated with the cardiac conduction system are affecting conduction. Whether the conduction system deletion model has a higher incidence of arrhythmia overall is ambiguous. No overt phenotype presented itself, and no gross changes to ECG wave morphology are noted.

In terms of contraction, the increased thickness of the ventricular walls, particularly the IVS, is notable. This septal thickening, coupled with the increased ejection fraction and left ventricular posterior wall thickness in the WT/Fl mice suggests the hearts are becoming hypercontractile. This change seems to affect both male and female mice. The IVS is the primary region in which MyBP-HL protein would have been deleted, and the fact that deletion from this small subset could induce septal hypertrophy warrants further investigation. These results again goes against the DCM phenotype seen in the constitutive null model [1] and also reinforce the idea that perhaps the DCM that develops in humans and in constitutive models is not directly from loss of MyBP-HL, but rather a second mechanism that is dysregulated after MyBP-HL loss.

More interesting is that the WT/Fl mice with this Contactin-2-Cre model are just as sick as the Fl/Fl, if not presenting a slightly worsened phenotype for some parameters. The WT/Fl Cre-positive group was the smallest of the three (*n = 7*), so perhaps that cohort is under sampled. It would be ideal to increase the number in that cohort more to see if the phenotype demonstrated by this data remains. The WT/Fl group being just as sick as the Fl/Fl is consistent with the constitutive model [1], demonstrating that even small perturbations in protein level are sufficient to cause some level of cardiac dysfunction.

Additionally, the females have an interesting and significantly increased E/A ratio after cardiac conduction system loss of *Mybphl* (p = 0.0003). The E/A value is a measure of the velocity of blood flow through the mitral valve during ventricular relaxation over atrial contraction. An increase in the E/A ratio could be a sign of increased left atrial pressure and may signify early atrial dysfunction [19]. It is possible this is the first indicator of atrial dysfunction in this model prior to extensive remodeling occurring. It is also notable that in both the conditional knock-down in adulthood and in this model, the female mice appeared to have the more notable functional changes, with worsened ejection fraction/fractional shortening in the Rosa-26-Cre(ERT2) mice and here with the significant E/A ratio increase, coinciding with a lack of physical changes to accompany them. Potentially, this represents an alternate disease progression in females where dysfunction presents in the absence of overt physical changes.

### MyBP-HL and Adrenergic Signaling

Both mouse models appear to respond to changes in adrenergic signaling. The conduction abnormalities are seen with isoproterenol and propranolol, each of which works via adrenergic receptors, although in opposite ways [14, 15]. Increased adrenergic response in heart disease is viewed as a compensatory mechanism and can cause sympathetic remodeling, including increased neuronal excitability, altered neurotransmitter production, and hyperinnervation or denervation of the cardiac tissue [20]. An increase in cardiac nerve growth factor occurs, which is caused by the greater adrenergic response, and this produces heterogeneous increases in nerve fiber density. This heterogeneity is proposed to lead to differing degrees of electrical remodeling within cardiomyocytes and thus a predisposition toward arrhythmias [20]. With loss of MyBP-HL from such a small subset of cells in the conduction system knock-out, the heterogeneous remodeling in very specific regions would make the differences between the recombined cells and the surrounding tissue even greater, potentially predisposing these regions to arrhythmia. The link between MyBP-HL and adrenergic signaling warrants further investigation of the specific pathway responsible for these changes.

There are several reasons why we may observe the hypercontractile phenotype in conditional knock-down mice. Potentially, the hearts becoming slightly hypercontractile is the first response to loss of the protein, then the hearts would become more dilated over time due to a secondary compensation mechanism. As stated previously, it has been shown that loss of MyBP-HL increases the linear relaxation phase in atrial sarcomeres, increasing cross-bridge cycling [3]. It may be the case that this increased rate of cycling, when chronic, causes the thin-walled atria to fatigue and leads to the atrial dilation seen in this model. In the case of the conduction system knock-down, perhaps the changes affect a small enough cell population that the mechanism that would normally confer DCM is not triggered. This would suggest that it is not the loss of MyBP-HL itself that causes DCM, but rather an alternative, potentially compensatory mechanism.

It would provide further clarity to age out a cohort of the conditional knock-down in adulthood mice and determine whether the hypercontractile phenotype turns into DCM or hypertrophic cardiomyopathy. Thus far, in the conditional knock-down model, changes in contraction are seen, but with DCM and HCM larger remodeling of the ventricle is typical, whether it be thinning or thickening of the left ventricular wall at rest. The conditional knock-down model has no changes in left ventricular diameter during diastole, so giving the hearts more time to remodel could further elucidate whether the hypercontraction is a temporary compensation that deteriorates to DCM or if the prolonged hypercontraction causes HCM.

A stressor such as trans-aortic constriction surgery or chronic isoproterenol exposure could also elicit a more pronounced phenotype. It is unexpected that an almost exclusively atrial phenotype causes ventricular dysfunction so soon after just partial loss of the protein in the adult conditional knock-down. Using a stressor would likely be more revealing in the cardiac conduction system knock-out than performing a longitudinal study. Based on the phenotype present after 18 weeks, the amount of time needed to develop a more apparent phenotype under stress-free conditions could be extensive. Trans-aortic constriction surgery would be effective for ventricular changes; however it is known that the surgery by itself will cause a pressure overload in the atria that would cause atrial dilation [21]. Whether the atrial dilation is significantly increased compared to the Cre-negative mice after surgery, and thus more than expected based on the ventricular remodeling, would be informative. For the conduction system knock-out mice, no atrial changes were noted, so performing the surgery could demonstrate whether loss of MyBP-HL in the atrial conduction system would confer quicker dilation and atrial dysfunction than the Cre-negative cohort.

It would also be beneficial to create a ventricular-specific deletion that could not only target the MyBP-HL positive cells within the VCS but also those adjacent to the VCS in the myocardium. While it does appear that there was sufficient deletion in the VCS, many puncta remained in the ventricles. This might give a more complete picture as to whether loss of MyBP-HL in all these cells would hinder the transition of the electrical signal in the heart from the Purkinje fibers to the myocardium.

The results from these models provide insights into primary atrial cardiomyopathies and their effect on the ventricles. Atrial dysfunction has often been viewed as secondary to ventricular disease, and it is rare that a link is studied between an atrial primary cardiomyopathy and ventricular dysfunction [22]. These results demonstrate that further study of MyBP-HL could uncover additional information regarding the way the atria and ventricles communicate and respond to stressors originating in the other chambers of the heart.

Based on these findings, people with loss-of-function MyBP-HL mutations are likely to have a higher risk of life-threatening arrhythmias and cardiovascular events. Patients with such mutations should be under consideration for receiving anti-arrhythmic medications and potentially ICD implantations at a younger age to mitigate these effects. As the discovery of this protein and its role in disease is relatively recent, *MYBPHL* does not appear on standard gene panel testing and little is known about the prevalence of mutations within various populations. Human data regarding the course of the disease is scarce because of this. Finding and then following more patients with these mutations would provide clarity regarding whether any contractile or conduction defects develop before the disease fully manifests and would allow further understanding as to how heart diseases progress in patients with *MYBPHL* mutations.

In conclusion, more work needs to be done to break down the exact mechanisms involved in disease progression from MyBP-HL knock-down or conduction-specific deletion. We can however conclude that even slight changes in the level of MyBP-HL create observable cardiac dysfunction, whether this occurs in adulthood or in only a small subpopulation of cells within the conduction system. This finding has implications for any missense mutations, which do not undergo nonsense mediated decay, but instead may affect the affinity of MyBP-HL to myosin or any other currently undiscovered binding partners. Therefore, MyBP-HL is essential for the heart as a whole to function normally, not just the atria.

## Abbreviations

(AF): atrial fibrillation
(AV): atrioventricular
(Cntn-2): cardiac myosin binding protein-C; contactin-2
(DCM): dilated cardiomyopathy
(HCM): hypertrophic cardiomyopathy
(IVS): interventricular septum
(LVID): left ventricular internal diameter
(LVPW): left ventricular posterior wall thickness
(MyBP-HL): myosin binding protein H-like
(MLC2a): myosin light chain 2a
(VCS): ventricular conduction system

